# Alpha-Amylase increases predict a freezing-like response and cortical alpha oscillations

**DOI:** 10.1101/2022.02.11.480062

**Authors:** Lisa Luther, Ole Jensen, Muriel A. Hagenaars

## Abstract

Individual differences in the reactivity of the sympathetic nervous system in response to stressful situations are thought to be an important predictor for psychological well-being and the focus of current scientific investigations. Here, we explored whether increased sympathetic nervous activity (SNA) was associated with reduced alpha power and with increased freezing-like behaviour (i.e., body sway) while watching threatening stimuli, reflecting enhanced attention. Salivary alpha-amylase (sAA) was used as a proxy for sympathetic nervous activity, which is elevated in stressful situations. A passive viewing task with affective pictures (unpleasant, neutral, pleasant) was carried out, and pre- and post- task sAA samples were taken. Oscillatory brain activity in the EEG and body sway were assessed simultaneously during the task. The results point to an increase in sAA being associated with reduced alpha power decrease to the unpleasant compared to the neutral pictures as well as increased freezing-like behaviour (i.e., reduction in body sway for unpleasant versus neutral pictures). It appears that an increase in SNA is linked to less attentional valence differentiation. Furthermore, our study corroborates findings from the animal literature in that the SNA increase is linked to a freezing-like response.

## Introduction

The individual response to stressful situations is accompanied by sympathetic nervous activity (SNA). SNA leads to enhanced secretion of the salivary enzyme alpha amylase (sAA). SAA has been established as a proxy for stress, based on animal as well as on human research. Through animal studies it has been established that sympathetic innervation of the salivary glands leads to high concentrations of sAA in saliva, whereas parasympathetic innervation leads to much lower concentrations of sAA in saliva (Anderson et al., 1984; Asking & Gjorstrup, 1987). The release of saliva is under control of neurotransmitters emitted by both sympathetic and parasympathetic neurons. Sympathetic neurons release noradrenaline (NA) which binds to saliva producing acinar cells in the salivary glands. Summarizing the cascade from sympathetic nervous activity (SNA) to an increase in sAA, Nater & Rohleder (2009) describe sympathetic neurons releasing NA which then binds to acinar cells in the salivary glands which in turn release saliva with high concentrations of sAA. Hence they describe sAA as biomarker for SNA. In human studies, stressful events have been related to increased sAA levels. For example, Ali and Pruessner (2012) found increased sAA levels after a Trier Social Stress Test. Takai and collegues (2004) found sAA increases during aversive video watching. Being exposed to unpleasant (and neutral) pictures has been shown to increase sAA levels as well: In an emotional rating task by Van Stegeren, Wolf, & Kindt (2008) participants were rating unpleasant and neutral pictures of the International Affective Picture System (IAPS; Lang, Bradley, Cuthbert., 2008) on emotionality. After the task (with a duration of approximately 30 minutes), sAA levels were increased. This is the only study to date to our knowledge relating affective picture viewing to an sAA increase.

Stress also modulates attention as reflected by specific brain activity patterns, as well as behavioural reactions. Regarding the former, experimental stressors such as aversive pictures have indeed been shown to readily draw attention (Balconi, Brambilla, & Falbo, 2009; Keil et al., 2001; Popov, Steffen, Weisz, Miller, & Rockstroh, 2012; Luther et al., in prep). Attentional processes in the brain are extensively studied, indicating that particularly alpha and gamma oscillations are modulated by attention (Foxe & Snyder, 2011; Jensen & Mazaheri, 2010; Klimesch et al., 2007). The alpha rhythm (8 – 14 Hz) has been shown to reflect inhibition of brain areas and hence reflect the inverse of attention, i.e. a decrease in alpha power reflects an increase in attention (Haegens et al., 2012; Klimesch et al., 2007; Okazaki et al., 2015), while gamma activity (30 – 120 Hz) reflects enhanced visual attention (Fries et al., 2001; Tallon-Baudry & Bertrand, 1999). Keil and collegues (2001) demonstrated a stronger decrease of alpha power for unpleasant compared to both pleasant and neutral pictures during right hemifield presentations. Consistently, Popov et al. (2012) found posterior alpha power to decrease less when passively watching neutral pictures compared to unpleasant pictures, indicating enhanced attention for unpleasant pictures. Is decreased alpha power as an indicator of increased attention towards unpleasant pictures related to physiological indicators of stress?

Aversive pictures have also been shown to elicit a behavioural stress response, i.e., freezing-like behaviour, defined as reduced body sway in response towards unpleasant compared to neutral or pleasant pictures (Azevedo et al., 2005; Hagenaars et al., 2012). In rodents, freezing was associated with increased adrenaline and noradrenaline (Nijsen et al., 2000), both indicating SNA and regulating attentional and behavioural threat responses. Importantly in humans, patients with posttraumatic stress disorder-relative to healthy controls-showed reduced freezing-like behaviour (Fragkaki et al., 2017). Stoffels and colleagues (2017) found bradycardia (also indicative of freezing) in response to aversive pictures in participants with low self-reported emotion avoidance but not in high emotion avoiding participants or patients with borderline personality disorder. Taken together, freezing-like behaviour seems to be healthy stress responding (including not avoiding emotional content) and (at least in animals) an increase in NE, indicative of SNA.

How are stress-level modulations related to threat-related attention and behavioural responses? The present study aims to shed light on whether increased SNA as indicative of an increased stress-level is linked to enhanced or decreased attention towards unpleasant pictures in humans. Similarly, we investigate whether physiological indicators of stress correlate with an increased freezing-like response, as it is in animals. Based on the above mentioned research, we will use sAA as proxy for SNA, EEG alpha and gamma power as proxy for attention, and body sway as proxy of a freezing-like response.

## Methods

### Participants

Fourty-four participants (18 male) were recruited from Radboud University Nijmegen, via a participation system, flyers, and personal contacts. Due to technical problems leading to incomplete data sets, data from four participants were excluded. Additionally, three more participants dropped out during the course of the experiment due to dizziness or feeling unwell, and 7 datasets contained too few trials after artifact rejection (< 10 trials in one condition), and of two participants there were no sAA measures. The mean age of the 28 remaining participants was 24.4 (*SD = 7.7*). The study was approved by the local ethics committee (Ethische Commissie Gedragswetenschappelijk onderzoek, ECG) and participants gave written informed consent prior to the experiment.

### Data Acquisition

#### sAA collection

Saliva was collected via a short straw into a little plastic cap. The tubes were put to freezing (ca. −80°C) at the end of the test day (Hansen et al., 2008). After data collection was finished, they were sent to the laboratory of Prof. Kirschbaum in Dresden for sAA analysis.

#### Apparatus

Stimuli were presented on a Samsung 2233SW screen with a refresh-rate of 60Hz or 120Hz and a resolution of 1920×1080 pixels. The screen was adjusted to eye-height at approximately 95 cm from the subject. A 64 channel EEG system (ActiCap64 system, Brain Products GmbH, Gilching, Germany) based on the extended 10-10 system was applied. The signals were referenced to linked mastoids and later re-referenced to a common reference. The data were sampled at 500 Hz, following band-pass filtering between 0.016 and 150 Hz. The recordings of the body sway took place on a custom build stabilometric square platform (50×50 cm; 19.7 inches) with a force sensor in each corner (distance between sensors: 42.5 cm; Niermann et al., 2015). The stabilometric data were acquired together with the EEG using four additional ExG channels.

### Procedure

Participants took part in a larger study, of which the third task was a passive viewing task of the IAPS pictures, before and after which salivary alpha-amylase was measured.

#### Stimuli

Each participant was presented 208 pictures from the International Affective Picture System (IAPS; Lang, Bradley, & Cuthbert, 2001) of which 52 were pleasant, 52 unpleasant, and 104 neutral pictures for passive viewing. Pictures were presented centrally with a visual angle of 8,9°. Pictures were selected based on the normative valence ratings provided by the IAPS manual (cut-off values: unpleasant < 3.2; neutral: 4 – 6; pleasant > 6.8). The valence ratings were further constrained not to differ more than 0.8 points between men’s and women’s ratings. The pictures were matched in terms of luminance and size with a custom written program. The same amount of pictures including humans (including the face) and non-humans was included in each category (double the amount in the neutral category, respectively).

#### Passive viewing task

Each picture appeared only once during the passive viewing task. Pictures were presented in 8 blocks consisting of 26 pictures. Each block contained 13 emotional pictures and 13 neutral pictures. This amounted to 4 blocks of pleasant and neutral pictures, and 4 blocks of unpleasant and neutral pictures. Thus, each block consisted half of neutral and half of arousing pictures (of either valence). The pictures and blocks were pseudo-randomized, being constrained to not show more than 2 blocks of one valence (pleasant or unpleasant) sequentially, and within each block not more than 4 pictures of one valence (neutral or pleasant, or neutral or unpleasant, respectively) in a row. Pictures were presented for 2 s with inter-trial intervals (ITI) of 1.5 – 2 s, leading to a total duration of around 1 min 36 s per block. A fixation-cross stayed on the screen and the pictures were presented around it (for details Luther et al. in prep). The blocks were separated by 20 s breaks. After 4 blocks followed a 3 min break allowing participants to sit down and rest their legs. During the break this cross turned into a countdown before resuming the picture presentation. Participants were asked to fixate on the cross while perceiving the picture around it. Furthermore, they were asked to blink after each picture in order to reduce blinking during stimulus presentation. Their posture was required to be relaxed with arms hanging alongside the body.

#### Alpha-Amylase collection

Saliva was collected directly before and immediately after the passive task, when participants had sat down. Time between the pre- and the post-measurement was approximately 25 minutes. See figure 1 for the study design

**Figure 1.**
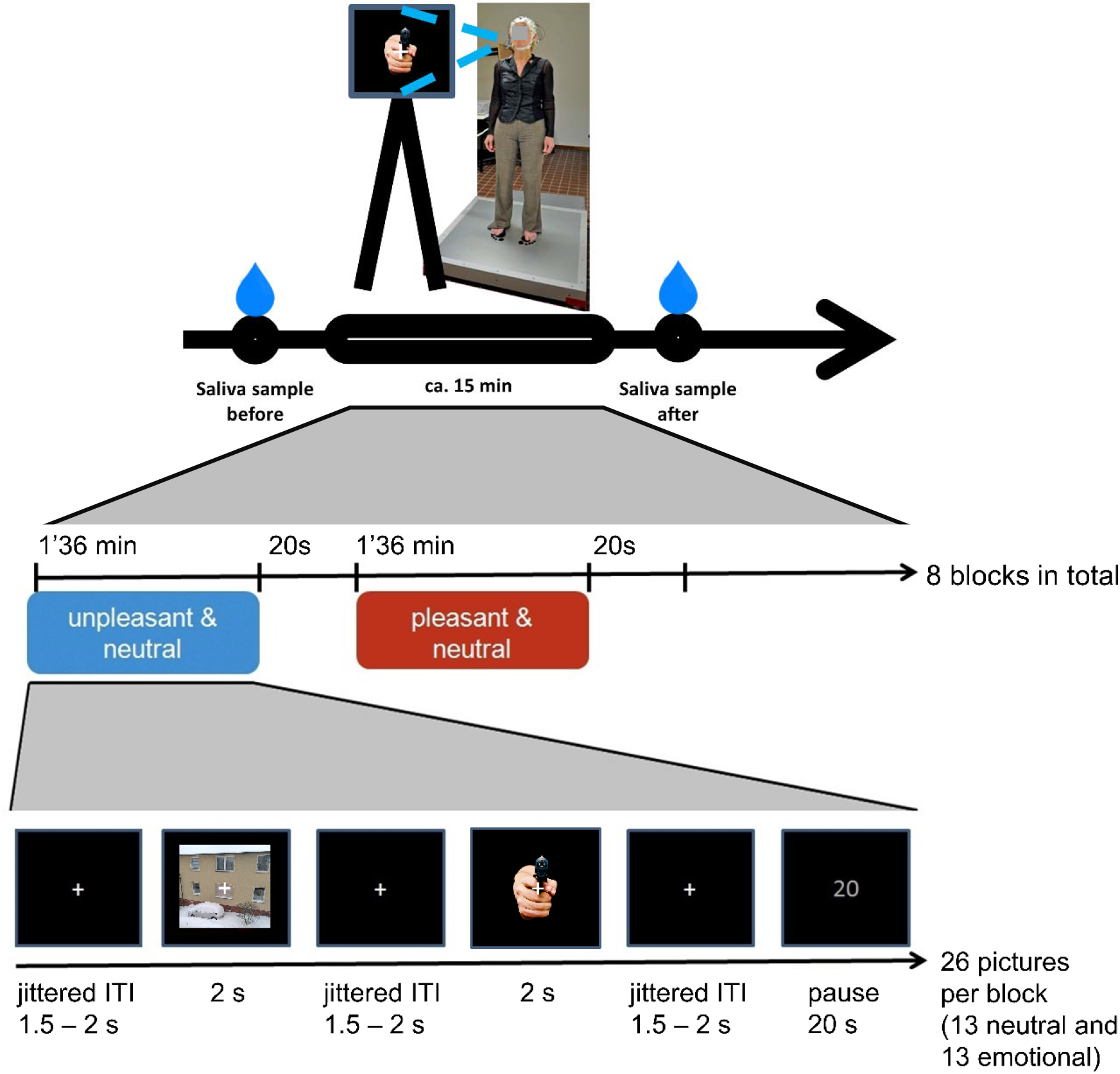
Experimental paradigm. Saliva samples were collected before and after a passive viewing task. During the passive viewing, participants were gazing at the fixation cross while perceiving the picture around the cross and EEG and body sway were recorded. From top to bottom: Top: study setup and time line of the experiment. Middle: time line of the passive viewing task in terms of conditions (blocks). Bottom: time line of the trial presentation within each block. Neutral pictures were presented among either pleasant or unpleasant pictures.

### Data analysis

#### Alpha-Amylase

We used a percent increase from pre- to post-passive viewing task (post minus pre divided by pre- plus post-measurement) for each subject. To account for expected changes in sAA due to diurnal rhythm (Figueiro & Rea, 2010; Marchand et al., 2016; Nater et al., 2007) we transformed the time of day at which subjects took part in the experiment into a sine-wave with an offset at 10am and peak at 4pm (trough at 4am and zero at 10pm, respectively) based on the cited literature. Thus, for each participant the pre- measurement time of day (e.g. 13.00h) was transformed with sin(2*pi/24*(timeofday −10)), and for the post-measurement 10.5 was subtracted leading to sin(2*pi/24*(timeofday −10.5)), respectively, since the time delay between pre- and post-measurement was about half an hour. This yielded a value between −1 and 1 indicating the expected phase of the sAA curve. Then the difference between the pre- and the post-value was taken (i.e. for each participant the slope on the sine wave was determined in terms of a single value) and that value was used as predictor for *expected* change in sAA. Participants took part between 12pm and 9pm of which 16 came before 4pm and 12 came after 4pm (and hence have a predicted increase (decrease, respectively) in sAA).

#### EEG sensor level analyses

EEG data were analyzed using MatLab (MathWorks Inc., 2014b) and the Matlab-based FieldTrip toolbox (Oostenveld, Fries, Maris, & Schoffelen, 2011). Artifacts were removed in a semi-automatic manner using visual inspection and selecting a threshold for each subject (Horschig et al., 2015). Trials containing ocular or muscle-artifacts were detected based on this threshold and excluded from analysis (on average 31%). Datasets with too many lost trials (less than 10 trials in one cell) were excluded from the EEG analysis.

Sensor level time-frequency analysis of power was performed using a 400 ms sliding time window (50ms steps) and a multi-taper approach (9 tapers; discrete prolate spheroidal sequences; DPSS, Percival & Walden, 1993) for the frequencies from 30 – 120 Hz; see e.g. Osipova et al., 2006).

In order to obtain a mean value for alpha and gamma power per condition per participant, we averaged power over the parietal sensors in the 0.5 – 1.5 s window and the 8 – 18 Hz (60 – 80 Hz, respectively) frequency range. This selection is based on the overall contrast for all conditions (F test) in Luther et al. (in prep.).

#### Stabilometric Platform

Offline, the data were low-pass filtered at 10 Hz. Body sway in anterior-posterior (AP) direction was calculated by summing the values of the two anterior sensors and of the two posterior sensors, respectively. Then standard deviation of movement in this direction (SD-AP) was calculated per trial over the stimulus period (0 – 2 s) and normalized to the baseline period (−1 – 0 s). Trials with a Z-score >4 (scores computed for each participant over all trials) were regarded as outliers and removed from analyses (on average 4 %). To normalize values across subjects to eliminate effects of body weight and height values were Z-scored, and then averaged per picture category (Hagenaars et al., 2012; Stins & Beek, 2007). Statistical analysis was conducted in SPSS 23 (SPSS, IBM Corporation). In order to obtain one value for body sway, we subtracted the mean SD-AP value of the neutral (pleasant, respectively) condition from the value of the unpleasant condition. The term freezing-like behaviour refers to the value after subtraction, i.e. it represents the condition difference.

#### Statistics

To test for a predictive value of an sAA increase on alpha and gamma power and body sway, a multiple regression with two independent variables (sAA percent increase, and sAA expected increase (controlling for effects of diurnal rhythm as explained above)) was applied on the different dependent variables (EEG alpha power, EEG gamma power, and body sway SD-AP), for the contrasts [unpleasant – neutral; *d*_(*un*)_] and [unpleasant – pleasant; *d*_(*up*)_] each.

## Results

Salivary Alpha-Amylase (sAA) was measured before and after a passive viewing task of unpleasant, neutral and pleasant pictures. EEG and body sway data were acquired throughout the task.

As described in Luther et al. (in prep.) we found robust modulations in the alpha and gamma band. In particular, we found a stronger alpha depression when unpleasant compared to pleasant pictures were presented (Luther et al.; in prep.). Similarly, we found a difference in the gamma band in which unpleasant pictures induced strong gamma power as compared to pleasant picture (Luther et al., in prep.). Also, we observed stronger gamma when unpleasant stimuli were compared to neutral stimuli (Luther et al., in prep.).

Based on those findings, we selected time, frequency and channels of interest (for exact parameters see methods section) over which we calculated the average power.

### Descriptives of the physiological measures of stress

The mean and standard deviation of the physiological measures (sAA, alpha and gamma power, body sway) are reported in Table 1.

**Table 1.**
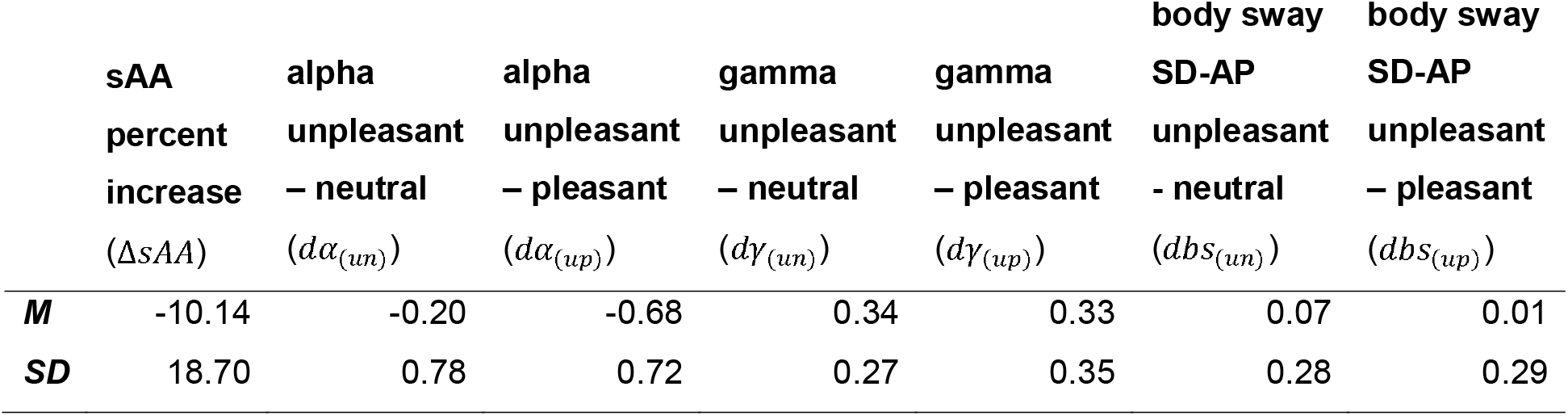
The unit of sAA is percent increase; EEG power is the *relative* alpha and gamma power (in μV) between the indicated conditions. SD-AP sway is the *relative* amount of variability in body movement between indicated conditions. Mean (M) and standard deviation (SD) of the relevant measures.

### Statistics

We used a general linear model to quantify changes in oscillatory activity in the context of affective or neutral stimuli. To test whether a change in sAA predicted emotion dependent EEG alpha power modulations, we conducted multiple regression analyses with the dependent variables being relative EEG alpha power ([unpleasant – neutral] and [unpleasant – pleasant]; *dα*_(*un*)_ and *dα*_(*up*)_), and percent change of sAA (pre-to post sAA; Δ*sAA*) being the predictor. In order to correct for the underlying diurnal rhythm of salivary alpha-amylase levels, we included the sAA’s expected increase due to time of day 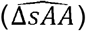 as second predictor (see methods section for details). This yielded the following six models (see equation 1 and 2 for alpha power; 3 and 4 for gamma power; 5 and 6 for body sway).

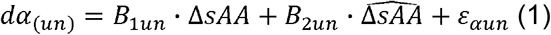

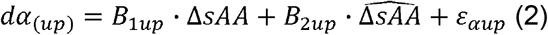

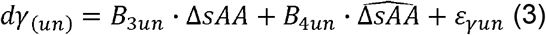

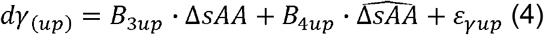

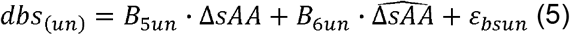

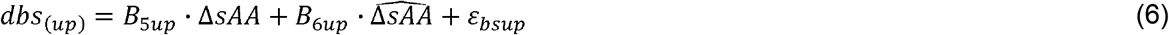

**Equations 1 to 6** | regression models for testing the relation between salivary alpha-amylase and posterior alpha power (gamma power and body sway in SD-AP direction, respectively). *dα*_(*un*)_: alpha power [unpleasant – neutral]; *dα*_(*up*)_: alpha power [unpleasant - pleasant]; *dγ*_(*un*)_: gamma power [unpleasant – neutral]; *dγ*_(*up*)_: gamma power [unpleasant - pleasant]; *dbs*_(*un*)_: body sway (SD-AP) [unpleasant – neutral]; *dbs*_(*up*)_: body sway (SD-AP) [unpleasant - pleasant]; *B*_*un*, *B*_*up*: beta weight for the subscripted contrast; Δ*sAA*: change in salivary alpha-amylase (pre- to post-task); 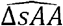: *expected* change in salivary alpha-amylase based on the time of day (see methods).

With the inclusion of an estimate of sAA time-of-day change next to the actual sAA change, it becomes important to check for collinearity. The most widely used metric for this is the variance inflation factor (VIF; Johnston et al., 2018). It is considered problematic if VIF > 2.5. Our VIF was 1.049, indicating no collinearity issues.

### Salivary Alpha-Amylase and EEG

#### EEG alpha power

Figure 2A shows the change in alpha power comparing unpleasant to neutral pictures in relation to percent change in sAA for the individual subjects. The core observation was that an increase in sAA was related to less alpha suppression towards unpleasant pictures compared to neutral pictures while a decrease in sAA showed the reverse. This was confirmed statistically since the overall model for EEG alpha power unpleasant vs. *neutral* pictures was significant (F(2,25) = 5.832, R^2^ = .318, p = .008). Both predictors (sAA measured increase and sAA expected increase due to time of day) contributed to the relative EEG alpha power difference between conditions (sAA percent increase: t = 2.123, β = .358, p = .044; see Figure 2A; sAA expected increase: t = −2.194, β = −.370, p = .038). This suggests that an increase in sAA entails slightly less attention towards the unpleasant compared to the neutral pictures.

**Figure 2.**
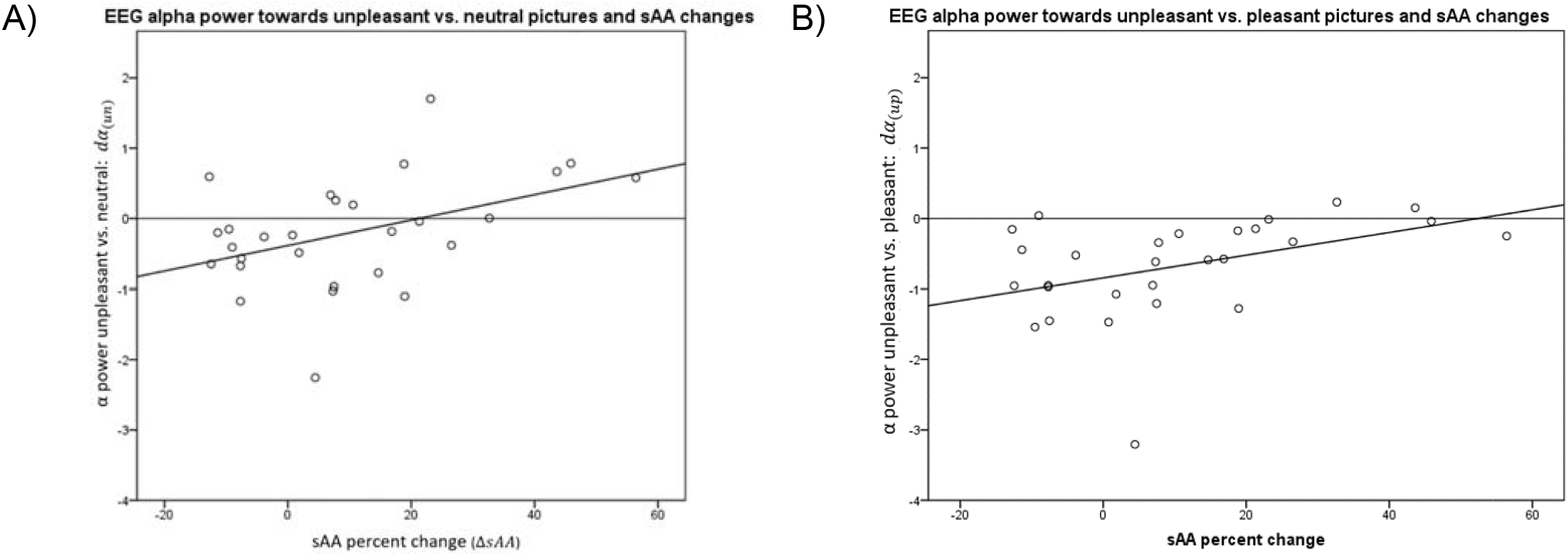
Salivary alpha-amylase (sAA) percent change from before to after the passive viewing of the IAPS pictures () and normalized EEG alpha power for (A) unpleasant pictures compared to neutral pictures (; t = 2.123, β = .358, p = .044) and (B) for unpleasant vs. pleasant pictures (; t = 2.033, β = .344, p = .053). With increasing sAA percent change (), subjects paid relatively less attention to unpleasant compared to neutral (and in a trend compared to pleasant) pictures as evidenced by a smaller EEG alpha power difference between conditions.

Figure 2B shows the change in alpha power comparing unpleasant to pleasant pictures in relation to changes in sAA. We observe that an sAA increase predicts an alpha modulation decrease while a decrease in sAA is related to a clear differentiation between the two conditions in terms of alpha modulation. Statistically, the overall model for EEG alpha power for unpleasant compared to *pleasant* pictures shows was significant: (F(2,25) = 5.658, R^2^ = .313, p = .009). Again, both predictors contributed though only close to significance for the percent increase (sAA percent increase: t = 2.033, β = .344, p = .053; see Figure 2B; sAA expected increase: t = −2.226, β = −.377, p = .035). Therefore, a relative decrease in alpha power towards unpleasant pictures compared to pleasant pictures is related to small changes in sAA.

In both models, the time-of-day-based expected sAA increase also predicts less attention towards unpleasant as compared to neutral and pleasant pictures, respectively.

#### EEG gamma power

In order to test for effects of sAA on EEG gamma power, we repeated the models with gamma power as the dependent variable. Here, we did not uncover significant effects for either contrast (unpleasant vs. *neutral*: F(2,25) = 1.298, R^2^ = .094, p = .291; sAA percent increase: t = −1.462, β = −.284, p = .156; sAA expected increase: t = 0.370, β = .072, p = .715; unpleasant vs. *pleasant*: F(2,25) = 0.578, R^2^ = .044, p = .568; sAA percent increase: t = −1.075, β = −.215, p = .293; sAA expected increase: t = −0.210, β = −.042, p = .836).

### Alpha-Amylase and body sway

We observed a change in body sway (SD-AP) comparing the unpleasant to the neutral condition in relation to changes in sAA for the individual subjects. The core observation was that increases in sAA were related to reduced body sway during unpleasant compared to neutral pictures. This was confirmed by a regression analysis in which the model for relative body sway between unpleasant and *neutral* pictures reaches F(2,25) = 2.418, p = .106, R^2^ = .162, with the single predictors contributing as follows: sAA percent increase: t = −2.187, β = −.408, p = .038 (see Figure 3); sAA expected increase: t = −0.796, β = .148, p = .434). Unpleasant compared to *pleasant* pictures reaches F(2,25) = 0.288, p = .752, R^2^ = .022, with the single factors contributing the following: sAA percent increase: t = 0.017, β = .003, p = .987; sAA expected increase: t = −0.746, β = −.151, p = .463. In both models, the expected change in sAA due to time-of-day is not related to body sway. Thus the increase in sAA related to less body sway towards unpleasant vs. neutral pictures, i.e., a freezing-like response seems to be related to an increase in sAA levels.

**Figure 3.**
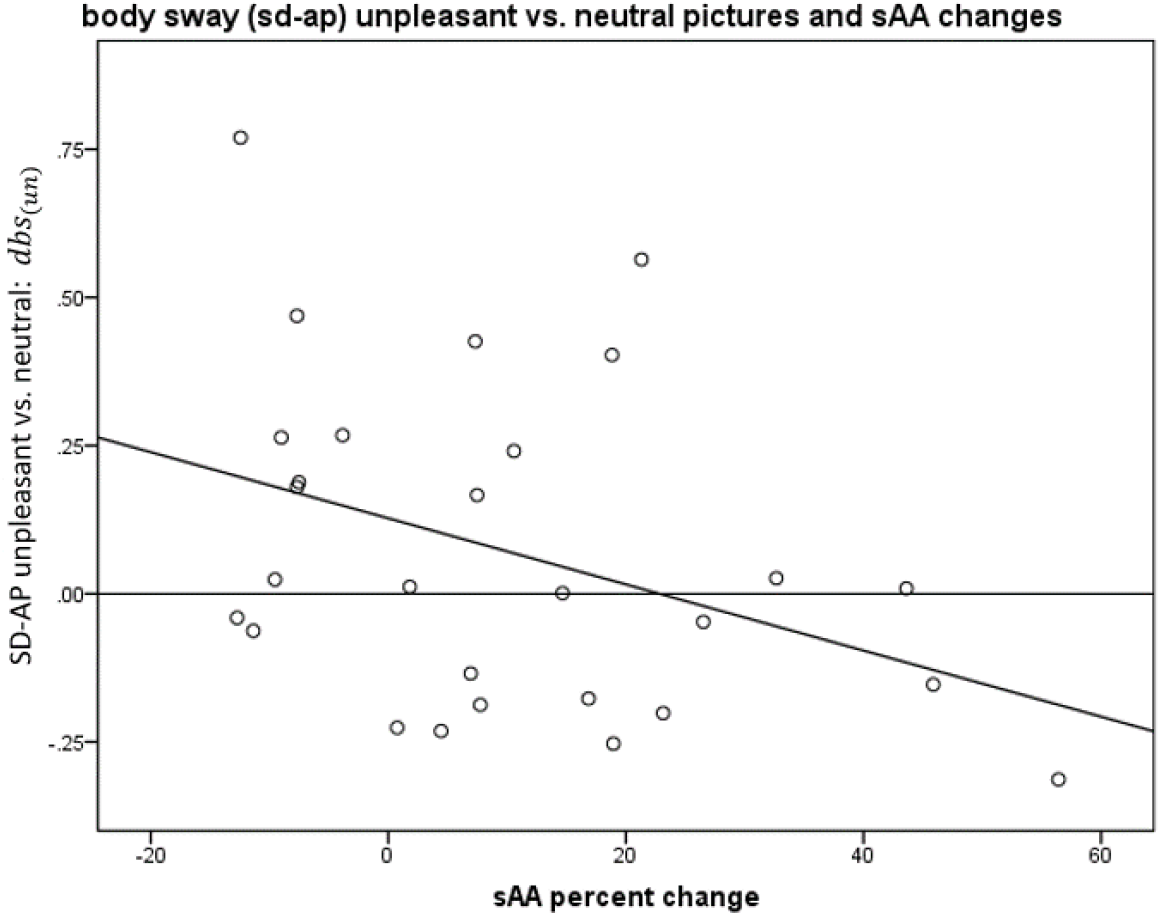
Salivary alpha-amylase (sAA) percent change from before to after the passive viewing of the IAPS pictures (Δ*sAA*) and body sway (SD-AP) for unpleasant compared to neutral pictures (*dbs*_(*un*)_; t = 2.187, β = .408, p = .038). With increased sAA levels body sway towards unpleasant pictures is decreased compared to neutral pictures (i.e. a freezing-like response is accompanied by an sAA increase)

## Discussion

In this study we tested the link between stress level modulation as indicated by pre- to post-task difference in salivary alpha-amylase concentration (sAA) as an indicator of sympathetic nervous activity (SNA), and stimulus induced modulations of visual alpha and gamma activity and freezing-like behaviour as indicated by body sway, towards unpleasant pictures as compared to neutral and pleasant pictures. The level of sAA increase/decrease per participant was related to the person’s average contrast [unpleasant – neutral] and [unpleasant – pleasant] for alpha activity and body sway respectively. Results indicate that an increase in pre to post sAA was related to less alpha suppression towards unpleasant pictures compared to neutral pictures (while a decrease in sAA showed the reverse). Similarly, we show a trend for increased differentiation in EEG alpha power between unpleasant and pleasant pictures with small changes in sAA; and smaller difference between unpleasant and pleasant pictures as sAA changes increase, indicating less discrimination in terms of attention with increased stress levels. For gamma activity this was not the case as no significant relationship with sAA levels was found. In addition, we found a strong trend for reduced sway for unpleasant compared to neutral (but not to pleasant pictures), i.e. an increased freezing-like response, as sAA levels increase. Note that while the reported effect were robust for the individual models they should be considered as strong trends when considering multiple comparisons.

### Decreased differentiation between conditions: hypervigilance and avoidance

In participants with higher stress response (higher sAA increases) to our task, we found two effects: a stronger freezing-like response, but also reduced differences in attention between unpleasant and neutral picture categories (as reflected by alpha modulations). The attention effects can be interpreted in two ways: participants might not pay particular attention to unpleasant pictures, or they pay inordinate attention to neutral and pleasant pictures. Such a state of indiscriminate reactivity could be due to general hypervigilance. Previous studies interpreted reduced behavioural discrimination (i.e., reduced difference in body sway) between unpleasant and neutral/pleasant stimuli as increased hypervigilance (Fragkaki et al., 2017; Hagenaars et al., 2012). Patients with PTSD display a general hypervigilance as a result of the traumatic event, and Fragkaki et al. (2017) found that this group did not show freezing in a task while the healthy control group did show a freezing-like response in the same task. Similarly, Hagenaars et al. (2012) found overall decreased body sway in participants with multiple aversive life events as compared to participants with no aversive life events, which they interpret as in line with PTSD phenomenology. Possibly, our alpha findings reflect similar processes as were reflected in these studies, but on another outcome measure (cortical alpha power instead of behavioural responses). Alternatively, the more stress-susceptible individuals may be adopting an avoidance strategy. In a study using target and distracting stimuli, Haegens et al. (2012) found that participants used increased alpha power to suppress distracting input in a somatosensory task, showing that increased alpha power can be used to suppress information processing. In this interpretation, stress-susceptible individuals in our study may be suppressing or avoiding the stimuli by attending less to them. In the current study it is not possible to distinguish between the two interpretations due to the lack of a pre-experiment baseline measurement. Future research should aim to incorporate this in order to distinguish between hypervigilance and decreased discrimination on baseline levels of attention. Either way, increased sAA seems to be related to both freezing-like behaviour and reduced attentional discrimination between unpleasant and neutral/pleasant stimuli. However, note that alpha and body sway were not found to be directly related in a previous investigation (Luther et al., in prep).

### Effect of time of measurement

In this study we see an indication of a freezing-like response linked to our proxy for SNA from pre- to post-task. Nagy et al. (2015) found that particularly the end of a cognitively stressful task coincides with an increase in sAA. Our study is different from the one by Nagy and colleges in that our task was emotionally stressful rather than cognitively. A more detailed focus on type of stress and sympathetic and parasympathetic patterns over the course of a study might be fruitful. The course of sAA concentrations could additionally be linked to body sway and EEG alpha and gamma power to understand the time course of physiological reactions to threat better.

### SNA and PNA (sympathetic and parasympathetic nervous activation)

In this study, we assume that sAA levels directly reflect SNA. However, it is possible that the relationship is more complex. Hagenaars et al. (2014) and Kozlowska et al., (2015) state that freezing is linked to a sympathetic and parasympathetic co-activation. In rats, it is this co-activation that produces the highest increase in sAA (Asking, 1985). However, whether this is true in humans as well has not yet been tested. Conducting a pharmacological study investigating whether at co-activation sAA concentration rates increase the most might be worthwhile, as sAA can be used as an relatively easy access measurement tool in studies on threat reactivity

### Effect of time of day on EEG alpha power

Our findings point to an interesting potential avenue of research, in that the time of day seems to be predictive of EEG alpha power in our condition contrasts, indicating that people attend more to unpleasant than to neutral/pleasant pictures in the evening and are less discriminative in terms of valence and arousal the in the morning.

### Limitations

We do find interesting results in terms of how sAA and body sway relate to the modulation of alpha power. While the reported individual models were statistically significant, the findings must be considered in the light of multiple comparisons given the 6 models. As such the reported effects should be considered strong trends. It would be of great interest to replicate the findings in future studies more statistical power. This could be achieved using a larger sample size and/or by better controlling for the variance of the dependent variables. For instance one limitation of our approach lies in the fact that the experiment was conducted at all times of day while sAA follows a circadian rhythm of its own. When limiting the time of day for measurements to a very confined range, sAA might yield more reliable results. We think that the mathematical correction we undertook eliminates most of this problem, but since it is impossible to perfectly catch the expected individual curves in a model, better practice would be to limit the time of day of participation.

### Future directions

With the current methodology, the causal link between emotion avoidance/hypervigilance and elevated stress biomarker levels effect cannot be directly established. Whether emotion avoidance ultimately leads to elevated enzymatic stress marker levels or whether people with elevated SNA use emotion avoidance or hypervigilance as (potentially suboptimal) strategy might be investigated in pharmacological studies by testing the effect of a direct manipulation of the enzymatic stress marker on alpha power or other measures of emotion avoidance. Additionally, it might be interesting to investigate the relationship between body sway and alpha/gamma power. Here, we assume the physiological measures to co-occur upon the detection of threat.

### Conclusion

From the current study we learned that increased SNA as measured by sAA levels is linked to less differentiation between unpleasant and neutral (a trend for pleasant pictures, respectively) in terms of EEG alpha power modulations, as well as an enhanced freezing-like response in terms of body sway (unpleasant vs. neutral). We interpret our findings as suggesting that less attentive differentiation between pictures of different valence (reflecting avoidance, hypervigilance, or indifference) and an enhanced freezing-like response are linked to a stronger increase in stress levels.

## References

Ali, N., & Pruessner, J. C. (2012). The salivary alpha amylase over cortisol ratio as a marker to assess dysregulations of the stress systems. Physiology and Behavior, 106(1), 65–72. https://doi.org/10.1016/j.physbeh.2011.10.003

Anderson, L. C., Garrett, J. R., Johnson, D. A., Kauffman, D. L., Keller, P. J., & Thulin, A. (1984). Influence of Circulating Catecholamines on Protein. J. Physiol., 352, 163–171.

Asking, B. (1985). Sympathetic stimulation of amylase secretion during a parasympathetic background activity in the rat parotid gland. Acta Physiologica Scandinavica, 124(4), 535–542. https://doi.org/10.1111/j.1748-1716.1985.tb00045.x

Asking, B., & Gjorstrup, P. (1987). Synthesis and secretion of amylase in the rat parotid gland following autonomic nerve stimulation in vivo. Acta Physiologica Scandinavica, 130(3), 439–445. https://doi.org/10.1111/j.1748-1716.1987.tb08160.x

Azevedo, T. M., Volchan, E., Imbiriba, L. A., Rodrigues, E. C., Oliveira, J. M., Oliveira, L. F., Lutterbach, L. G., & Vargas, C. D. (2005). A freezing-like posture to pictures of mutilation. Psychophysiology, 42(3), 255–260. https://doi.org/10.1111/j.1469-8986.2005.00287.x

Balconi, M., Brambilla, E., & Falbo, L. (2009). Appetitive vs. defensive responses to emotional cues. Autonomic measures and brain oscillation modulation. Brain Research, 1296, 72–84. https://doi.org/10.1016/j.brainres.2009.08.056

Figueiro, M. G., & Rea, M. S. (2010). The Effects of Red and Blue Lights on Circadian Variations in Cortisol, Alpha Amylase, and Melatonin. International Journal of Endocrinology, 2010, 1–9. https://doi.org/10.1155/2010/829351

Foxe, J. J., & Snyder, A. C. (2011). The role of alpha-band brain oscillations as a sensory suppression mechanism during selective attention. Frontiers in Psychology, 2(JUL), 1–13. https://doi.org/10.3389/fpsyg.2011.00154

Fragkaki, I., Roelofs, K., Stins, J., Jongedijk, R. A., & Hagenaars, M. A. (2017). Reduced freezing in posttraumatic stress disorder patients while watching affective pictures. Frontiers in Psychiatry, 8(MAR), 1–9. https://doi.org/10.3389/fpsyt.2017.00039

Fries, P., Reynolds, J. H., Rorie, A. E., & Desimone, R. (2001). Modulation of oscillatory neuronal synchronization by selective visual attention. Science (New York, N.Y.), 291(5508), 1560–1563. https://doi.org/10.1126/science.291.5508.1560

Haegens, S., Luther, L., & Jensen, O. (2012). Somatosensory Anticipatory Alpha Activity Increases to Suppress Distracting Input. Journal of Cognitive Neuroscience, 24(3), 677–685. https://doi.org/10.1162/jocn_a_00164

Hagenaars, M. A., Oitzl, M., & Roelofs, K. (2014). Updating freeze: Aligning animal and human research. Neuroscience and Biobehavioral Reviews, 47, 165–176. https://doi.org/10.1016/j.neubiorev.2014.07.021

Hagenaars, M.A., Stins, J. F., & Roelofs, K. (2012). Aversive life events enhance human freezing responses. Journal of Experimental Psychology: General, 141(1), 98–105. https://doi.org/10.1037/a0024211

Hansen, Å. M., Garde, A. H., & Persson, R. (2008). Sources of biological and methodological variation in salivary cortisol and their impact on measurement among healthy adults: A review. Scandinavian Journal of Clinical and Laboratory Investigation, 68(6), 448–458. https://doi.org/10.1080/00365510701819127

Horschig, J. M., Smolders, R., Bonnefond, M., Schoffelen, J. M., Van Den Munckhof, P., Schuurman, P. R., Cools, R., Denys, D., & Jensen, O. (2015). Directed communication between nucleus accumbens and neocortex in humans is differentially supported by synchronization in the theta and alpha band. PLoS ONE, 10(9), 1–20. https://doi.org/10.1371/journal.pone.0138685

Jensen, O., & Mazaheri, A. (2010). Shaping functional architecture by oscillatory alpha activity: gating by inhibition. Frontiers in Human Neuroscience, 4, 186. https://doi.org/10.3389/fnhum.2010.00186

Johnston, R., Jones, K., & Manley, D. (2018). Confounding and collinearity in regression analysis: a cautionary tale and an alternative procedure, illustrated by studies of British voting behaviour. Qual Quant. https://doi.org/10.1007/s11135-017-0584-6

Keil, A., Mueller, M. M., Gruber, T., Wienbruch, C., Stolarova, M., & Elbert, T. (2001). Effects of emotional arousal in the cerebral hemispheres ◻: a study of oscillatory brain activity and event-related potentials. 112, 2057–2068.

Klimesch, W., Sauseng, P., & Hanslmayr, S. (2007). EEG alpha oscillations: The inhibition-timing hypothesis. Brain Research Reviews, 53(1), 63–88. https://doi.org/10.1016/j.brainresrev.2006.06.003

Kozlowska, K., Walker, P., McLean, L., & Carrive, P. (2015). Fear and the Defense Cascade: Clinical Implications and Management. Harvard Review of Psychiatry, 23(4), 263–287. https://doi.org/10.1097/HRP.0000000000000065

Lang, P. J., Bradley, M. M., & Cuthbert, B. N. (2008). International affective picture system (IAPS): Affective ratings of pictures and instruction manual. Technical Report A-8.

Luther, L., Horschig, J.M., van Peer, J.M., Roelofs, K., Jensen, O., Hagenaars, M.A. (in prep.). Oscillatory Brain Responses to Emotional Stimuli are Effects Related to Events rather than States.

Marchand, A., Juster, R. P., Lupien, S. J., & Durand, P. (2016). Psychosocial determinants of diurnal alpha-amylase among healthy Quebec workers. Psychoneuroendocrinology, 66, 65–74. https://doi.org/10.1016/j.psyneuen.2016.01.005

Nagy, T., van Lien, R., Willemsen, G., Proctor, G., Efting, M., Fülöp, M., Bárdos, G., Veerman, E. C. I., & Bosch, J. A. (2015). A fluid response: Alpha-amylase reactions to acute laboratory stress are related to sample timing and saliva flow rate. Biological Psychology, 109, 111–119. https://doi.org/10.1016/j.biopsycho.2015.04.012

Nater, U. M., & Rohleder, N. (2009). Salivary alpha-amylase as a non-invasive biomarker for the sympathetic nervous system: Current state of research. Psychoneuroendocrinology, 34(4), 486–496. https://doi.org/10.1016/j.psyneuen.2009.01.014

Nater, U. M., Rohleder, N., Schlotz, W., Ehlert, U., & Kirschbaum, C. (2007). Determinants of the diurnal course of salivary alpha-amylase. Psychoneuroendocrinology, 32(4), 392–401. https://doi.org/10.1016/j.psyneuen.2007.02.007

Niermann, H. C., Ly, V., Smeekens, S., Figner, B., Riksen-Walraven, J. M., & Roelofs, K. (2015). Infant attachment predicts bodily freezing in adolescence: evidence from a prospective longitudinal study. Front Behav Neurosci, 9(October), 263. https://doi.org/10.3389/fnbeh.2015.00263

Nijsen, M. J. M. A., Croiset, G., Diamant, M., Stam, R., Kamphuis, P. J. G. H., Bruijnzeel, A., De Wied, D., & Wiegant, V. M. (2000). Endogenous corticotropin-releasing hormone inhibits conditioned-fear-induced vagal activation in the rat. European Journal of Pharmacology, 389(1), 89–98. https://doi.org/10.1016/S0014-2999(99)00870-5

Okazaki, Y. O., Horschig, J. M., Luther, L., Oostenveld, R., Murakami, I., & Jensen, O. (2015). Real-time MEG neurofeedback training of posterior alpha activity modulates subsequent visual detection performance. NeuroImage, 107, 323–332. https://doi.org/10.1016/j.neuroimage.2014.12.014

Oostenveld, R., Fries, P., Maris, E., & Schoffelen, J. M. (2011). FieldTrip: Open source software for advanced analysis of MEG, EEG, and invasive electrophysiological data. Computational Intelligence and Neuroscience. https://doi.org/10.1155/2011/156869

Osipova, D., Takashima, A., Oostenveld, R., Fernandez, G., Maris, E., & Jensen, O. (2006). Theta and Gamma Oscillations Predict Encoding and Retrieval of Declarative Memory. Journal of Neuroscience, 26(28), 7523–7531. https://doi.org/10.1523/JNEUROSCI.1948-06.2006

Popov, T., Steffen, A., Weisz, N., Miller, G. A., & Rockstroh, B. (2012). Cross-frequency dynamics of neuromagnetic oscillatory activity: Two mechanisms of emotion regulation. Psychophysiology, 49(12), 1545–1557. https://doi.org/10.1111/j.1469-8986.2012.01484.x

Stins, J. F., & Beek, P. J. (2007). Effects of affective picture viewing on postural control in healthy male subjects. BMC Neuroscience, 83(8), 202–210. https://doi.org/10.1186/1471-2202-8-83

Stoffels, M., Nijs, M., Spinhoven, P., Mesbah, R., & Hagenaars, M. A. (2017). Emotion avoidance and fear bradycardia in patients with borderline personality disorder and healthy controls. Journal of Behavior Therapy and Experimental Psychiatry, 57, 6–13. https://doi.org/10.1016/j.jbtep.2017.02.001

Takai, N., Yamaguchi, M., Aragaki, T., Eto, K., Uchihashi, K., & Nishikawa, Y. (2004). Effect of psychological stress on the salivary cortisol and amylase levels in healthy young adults. Archives of Oral Biology, 49(12), 963–968. https://doi.org/10.1016/j.archoralbio.2004.06.007

Tallon-Baudry, C., & Bertrand, O. (1999). Oscillatory gamma activity in humans and its role in object representation. Trends in Cognitive Sciences, 3(4), 151–162. https://doi.org/10.1016/s1364-6613(99)01299-1

van Stegeren, A. H., Wolf, O. T., & Kindt, M. (2008). Salivary alpha amylase and cortisol responses to different stress tasks: Impact of sex. International Journal of Psychophysiology, 69(1), 33–40. https://doi.org/10.1016/j.ijpsycho.2008.02.008

